# Quantifying network behavior in the rat prefrontal cortex: a reproducibility crisis

**DOI:** 10.1101/2023.05.16.541018

**Authors:** Congzhou M. Sha, Jian Wang, Richard B. Mailman, Yang Yang, Nikolay V. Dokholyan

## Abstract

The question of how consciousness and behavior arise from neural activity is fundamental to understanding the brain, and to improving the diagnosis and treatment of neurological and psychiatric disorders. There is significant murine and primate literature on how behavior is related to the electrophysiological activity of the medial prefrontal cortex and its role in working memory processes such as planning and decision-making. Existing experimental designs, however, have insufficient statistical power to unravel the complex processes of the prefrontal cortex. We therefore examined the theoretical limitations of such experiments, providing concrete guidelines for robust and reproducible science. We piloted the use of dynamic time warping and associated statistical tests to data from neuron spike trains and local field potentials, to quantify neural network synchronicity and correlate neuroelectrophysiology with rat behavior. Our results indicate the statistical limitations of existing data, making meaningful comparison between dynamic time warping with traditional Fourier and wavelet analysis currently impossible until larger and cleaner datasets are available.

**Significance Statement:** The prefrontal cortex is important in decision-making, yet no robust method currently exists to correlate neuron firing in the PFC to behavior. We argue that existing experimental designs are ill-suited to addressing these scientific questions, and we propose a potential method using dynamic time warping to analyze PFC neural electrical activity. We conclude that careful curation of experimental controls is needed to separate true neural signals from noise accurately.

## Introduction

Hodgkin and Huxley’s model of neuron action potentials derived from experiments with the squid giant axon^1^ was the seminal event in neurophysiology^2–4^. The central dogma of modern neuroscience is that neuron electrochemical activity and connectivity at the microscopic level can provide a clear understanding of complex behaviors. Thus, measuring electrical signals can be sufficient to integrate the chemical signaling and activity of neural networks.

The visual cortex and its relationship to visual processing is one of the better-understood neural networks because there are significant spatial correlations in signals that result from local connections between neurons. These spatial autocorrelations result in a high signal-to-noise ratio due to the strong intensity of the electric field produced by local synchronized neuron firing. Additionally, the neuroelectrophysiology of individual cells in the visual pathway has allowed mathematical modeling of the performance of specific neurons^5,6^. The spatial autocorrelations of visual neural activity are sufficiently high that low-resolution, indirect measurements of neuron activity, such as local blood flow^7–9^ and extracranial electrodes^10^, are sufficient to reconstruct perceived images. The organization of the visual cortex may be explained by the fact that images projected on the retina are spatially correlated, and therefore biological neural networks have adapted to take advantage of these correlations.

From an evolutionary perspective, the visual cortex is an ancient structure, and therefore the neural networks involved in visual processing have optimized over time. In contrast, the prefrontal cortex as part of the neocortex is one of the least developed brain regions, especially through evolutionary analysis of its size in mammals^11,12^. Unlike the visual cortex, the prefrontal cortex does not seem to possess highly local and regular spatial organization^13^. Because of the lack of regular spatial organization of the prefrontal cortex and the consequent low intensity of electrical signals, electrophysiology is one of the only methods with adequate sensitivity to probe the fine structure of the prefrontal neural network. Unfortunately, such measurements are highly invasive, and experiments are only feasible on laboratory animals such as rats^14–16^. To find the relationship between neuron firing and behavior, methods are required to analyze these complex electrical measurements.

The characteristics of this neuron prefrontal network constrain the firing patterns of neurons. Therefore, methods are required to capture key network characteristics from recorded spike trains and local field potentials (LFPs). Our hypothesis is that these network characteristics are embodied by the level of synchronicity of neuron firing: neurons that work in concert will have spike trains and/or LFPs that are highly correlated or anti-correlated, whereas neurons that process independent data will not be correlated.

When analyzing complex data such as neuroelectrophysiology tracings, the major statistical concern is bias and overfitting^17^. Although large amounts of data are collected in these experiments, the degrees of freedom are severely restricted by the number of animals and recordings collected^18^. When researchers choose a statistical model for their data, certain assumptions are made about the structure of the data. Commonly, a class of statistical models is chosen with hyperparameters tuned to best fit the data. The more hyperparameters that must be tuned, the fewer degrees of freedom remain for the subsequent goodness-of-fit tests. In this work, we sought to maximize the degrees of freedom remaining after modeling to address our scientific questions. Schematically, we must allocate the degrees of freedom in the data to the modelling we perform and the scientific questions we seek to answer:

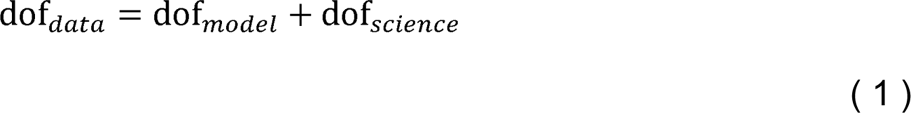

The more degrees of freedom we can devote to science, the more confident we can be in our statistical tests.

Therefore, we chose to investigate neuron synchronicity using the dynamic time warping (DTW) method^19^. DTW is an efficient, non-parametric approach to determine the best alignment of two time series, such that the overall shapes of the time series are matched. Compared to classical Fourier and wavelet analysis^14,15^, DTW gives a global alignment of the data that is robust to local variations in timing. We predicted that DTW could discern if two neurons fire synchronously or asynchronously. In contrast, local timing variations in low frequency signals such as firing rate and voltage lead to destructive interference when performing the Fourier or wavelet transform, decreasing the signal-to-noise ratio. In practice, these techniques require the tuning of various hyperparameters to smooth and denoise the data. Examples include a window size for the spike-triggered LFP average^14^, cutoff frequencies for band-pass filters^14^, various bootstrapping techniques^14^, and choice of kernel for support vector machines^15^. In contrast, traditional DTW uses no additional parameters to yield a dissimilarity index for each pair of spike trains or LFP tracings, therefore retaining more degrees of freedom for answering statistical questions.

Thus, we analyzed electrophysiology data collected from rats performing the T-maze task, a task that evaluates working memory. We use previously published spike train and LFP recordings taken from the rat medial prefrontal cortex (mPFC)^14–16^, along with previously unpublished data from three additional rats (rats A, B, and C) recorded using published methods^16^. We investigated if DTW would be useful in elucidating the connection between neural activity and behavior (Results), and we examined the underlying assumptions of neuroelectrophysiology experiments (Discussion).

## Results

Our theoretical arguments for the low statistical power of existing experimental designs are presented in the Discussion and Methods (*Reproducibility of electrode recordings*). The data from the three published studies (Table 1) had rat neuron spike trains and/or LFPs recorded as the rats performed in T-maze (Figure 1-A). The exact experimental details for the three studies differed slightly^14–16^, but their combined scientific goal was to analyze the link between neuron recordings and working memory performance in the T-maze task. We focused on the four-second window of time centered on the moment at which the rat leaves the T-intersection (decision box in Figure 1-A), presumably making its choice. This window contained two seconds of pre-decision neural activity (e.g., cognition and decision-making process) and two seconds of post-decision neural activity (e.g., evaluation of reward or lack of reward). Since the experimental designs differed among the studies (e.g., the rat either traversed the T-maze continuously^14,15^ or was picked up by the researcher in between traversals^16^), we used this limited window of time for our analysis to limit the effects of the design, constituting one of the only hyperparameters chosen in our analysis. Our hypothesis was that neural activity was affected significantly by the independent variables of (1) timing of the recording relative to the choice, (2) correctness of the choice, and (3) sampled neurons (Table 2). Null and alternative hypotheses for each statistical test we performed are included in the Supplemental Tables.

**Figure 1:**
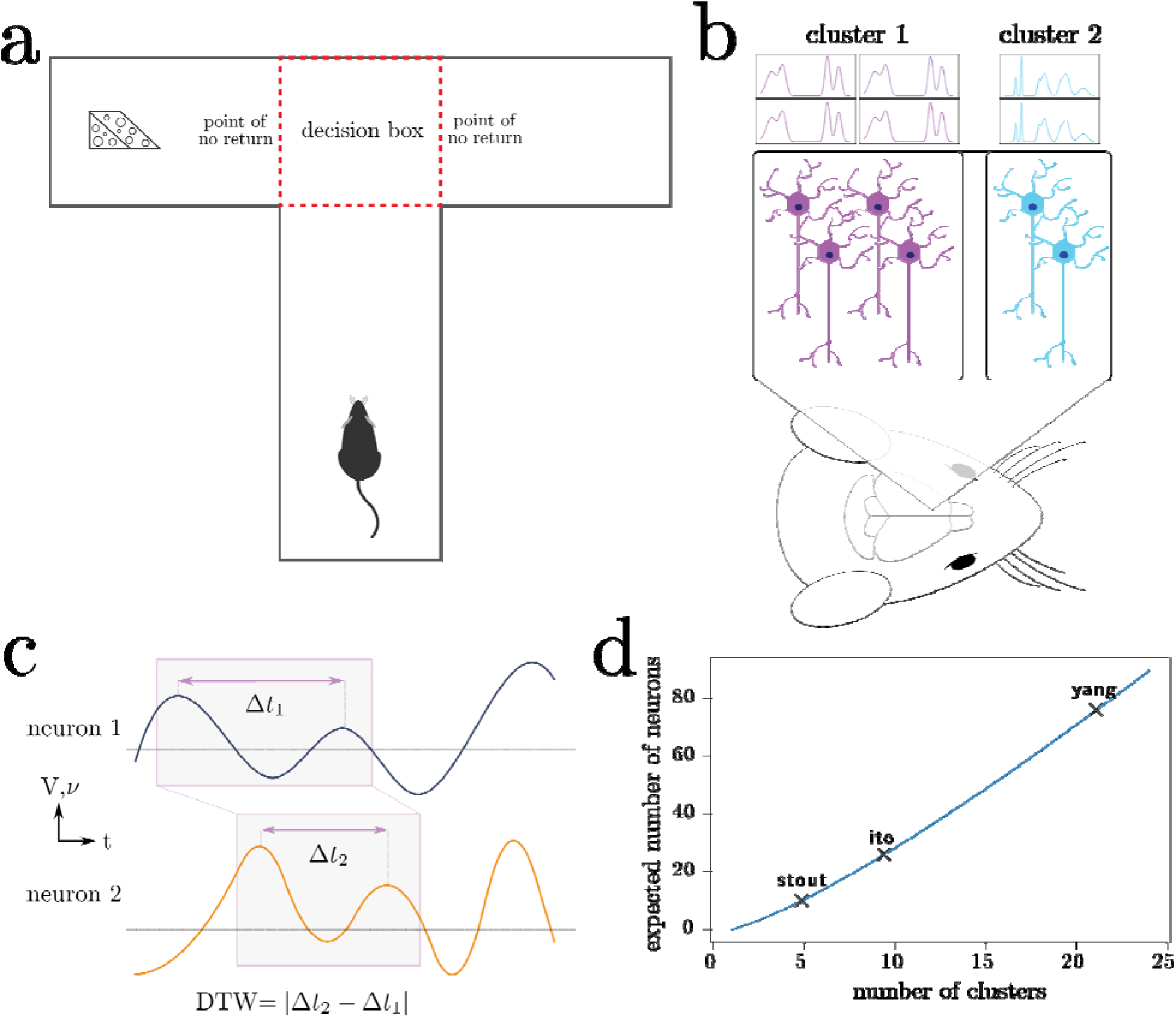
Outline of experiments, our working model, and data analysis. (a) Rats complete the T-maze task, in which the reward switches locations after each completion of the task. This task tests the working memory and decision-making capacity of the rat. (b) Neurons in the medial prefrontal cortex are recorded using intracranial electrodes, resulting in the measurement of specific clusters of neurons. (c) Dynamic time warping computes an optimal alignment of neuron spike trains and local field potentials, producing a single number characterizing the synchronicity of the neurons. (d) Depending on the number of distinct neuron clusters in the region of interest, a certain number of neurons must be sampled to produce a representative sample of all the clusters; in combinatorics, this is known as the coupon collector problem. We plot the number of clusters that the maximum number of neurons recorded in each of the three studies can theoretically discern.

**Table 1:** Characteristics of the three studies analyzed in this work.

|  | Ito<sup>14</sup> | Stout<sup>15</sup> | Yang<sup>16</sup> |
| --- | --- | --- | --- |
| Year of publication | 2018 | 2020 | 2022 |
| Number of rats | 3 | 5 | 9 |
| Total number of sessions<br>(including sessions in which only<br>one neuron was recorded) | 13 | 45 | 13 |
| Sessions excluded (sessions in<br>which only a single neuron was<br>recorded) | 0 | 9 | 0 |
| Number of trials (excluding single-<br>neuron recordings) | 790 | 1508 | 472 |
| Mean neurons recorded (s.d.) | 17.56 (4.24) | 3.31 (2.21) | 31.8 (28.55) |
| Range of neurons recorded | [10, 26] | [1, 10] | [5, 76] |
| Fixed electrode | No | No | Yes |
| Spatial metadata | Yes | Yes | Yes |

**Table 2:** Listing of categorical variables used for stratification/grouping in our analyses.

| Categorical variable | Values | Interpretation |
| --- | --- | --- |
| timing | before, after | Data from before vs. after the rat left the |
|  |  | T-intersection |
| correctness | true, false | True if rat chose the correct (i.e. alternate) branch of the T-maze during the task |
| study | Ito, Stout, Yang | The study to which the rat belongs |
| rat | 17 total animals | The rat identity |
| session | 62 total sessions<br>(with more than 1<br>neuron recorded) | The rat identity and the date of recording |

We used the DTW method on the neuron spike trains from all three studies and from the local field potentials for Ito et al.^14^ and Yang et al.^16^ (Figure 1-B & 1-C). Since DTW provides a numerical measurement of dissimilarity between firing of neurons, we converted the resulting DTW matrices into undirected, unweighted graphs (Supplemental Methods). For each set of neuron spike trains and/or LFPs, we quantified the connectivity of the resulting DTW graph by a single number, *d*_crit_. To evaluate for the presence of differences among sets of experiments, we used non-parametric statistical tests (Kruskal-Wallis, Kolmogorov-Smirnov, Mantel, and Boschloo exact tests) to mitigate both our lack of knowledge of the true underlying probability distributions and the small sample sizes. These statistical tests and the calculation of *d*_crit_ for a given DTW matrix are described in detail in the Supplemental Methods. We visualized *d*_crit_ as a function of our independent variables in Figure 2 (spike trains) and Supplemental Figure 1 (LFPs).

**Figure 2:**
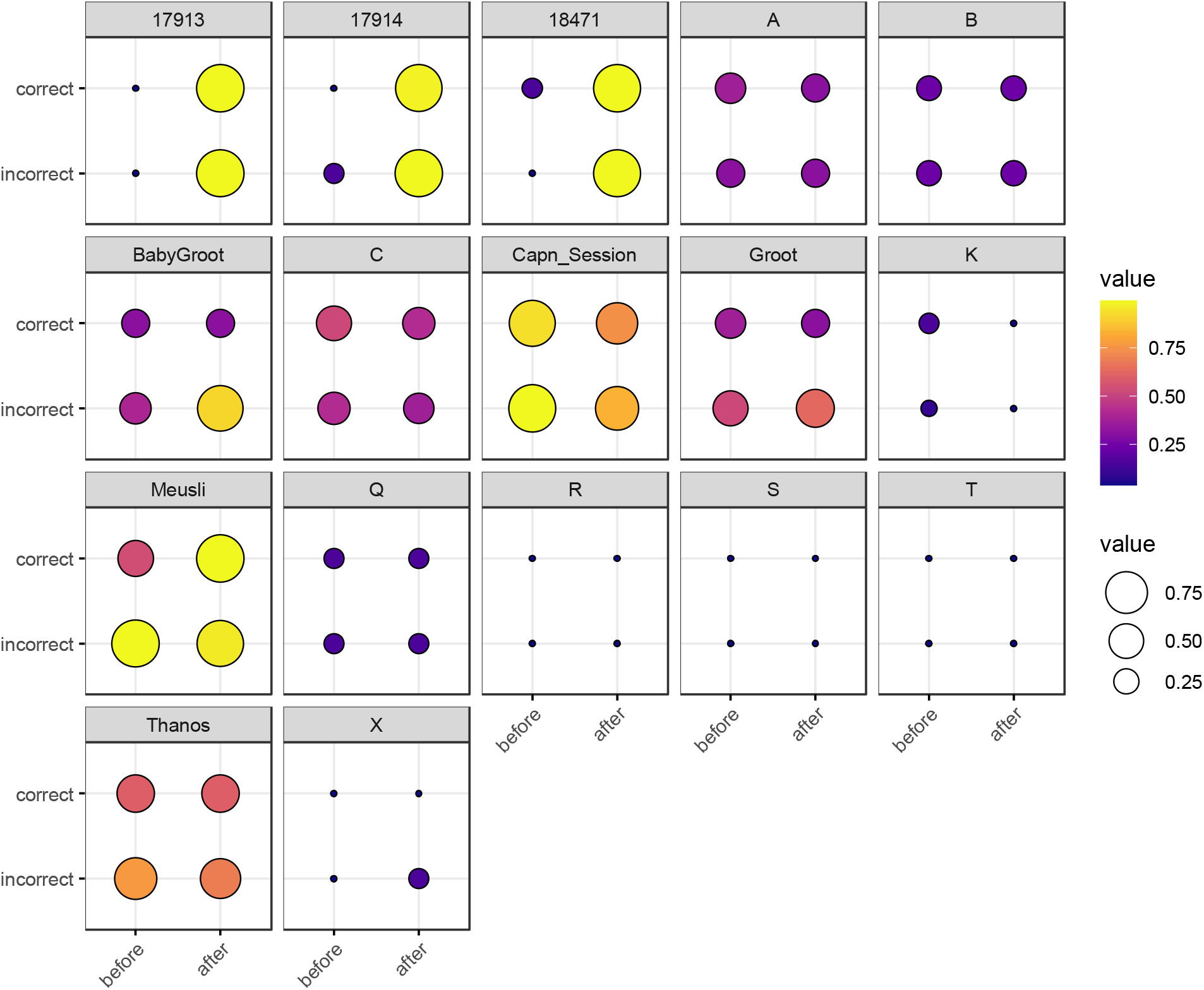
Balloon plot of neuron spike train *d_crit_*, grouped by rat, correctness of T-arm choice, and timing with respect to T-arm choice. Each box represents a single rat. The x-axis represents timing (before vs. after the rat visits the T-maze intersection) and the y-axis represents correctness of T-arm taking (True vs. False). The size and color of each marker represents log_10_*d_crit_*. For rats with multiple trials, we took the mean of log_10_*d_crit_*. Rats 17913, 17914, and 18471 are from Ito^14^ (2018); single letter rats are from Yang^16^ (2022); the remaining five rats are from Stout^15^ (2020). The most visually striking difference is in the Ito^14^ rats, in which there is a stark contrast for spike trains recorded before vs. after making a choice.

We defined a ***trial*** as a single traversal of the T-maze by a rat. We defined a ***session*** as all trials recorded on a single day for a single rat. In the Ito^14^ and Stout^15^ studies, electrodes were adjusted between sessions, and, therefore, a different set of neurons were recorded between sessions. We set the level of statistical significance (false positive rate) to α = 0.05.

### Neural network firing is consistent across a single session

Our first question was whether in analyzing *d*_crit_, could we pool trials from a single session? The null hypothesis was that during a single session, we may assume that the rat basal activity is unchanged between recordings in the same session, as measured by *d*_crit_, The alternative hypothesis is that specific factors affect *d*_crit_ between trials in the same session, such as the categorical variables in Table 2.

First, we analyzed the normality of *d*_crit_ with the Shapiro-Wilk test (Supplemental Tables 1 and 2). For neuron spike trains, we found that in 96.7% (60 out of 62) of sessions, the samples were not normally-distributed (*p* < 0.05), and for LFPs, we found that 80.0% (20 out of 25) were not normally-distributed. Comparing spike trains to LFP with Boschloo’s exact test, these fractions were not significantly different (p=0.615). We performed subgroup analysis on the categorical variables (1) correctness and (2) timing and found that this proportion was unaffected by further stratification by these variables. Therefore, we concluded that the Kruskal-Wallis non-parametric test was more appropriate than classical one-way analysis of variance (ANOVA), due to violation of the assumption of normally-distributed data and due to the limited number of trials per session.

Only 4.8% (3 out of 62) of the spike train sessions displayed significant (Kruskal-Wallis *p* < 0.05) trial-to-trial variance of *d*_crit_, regardless of stratification by correctness and/or timing; the corresponding proportion for LFPs was 12% (3 out of 25). For completeness, we also performed ANOVA and found that 11.2% (7 out of 62) of the spike train sessions displayed significant trial-to-trial variance, regardless of stratification; the corresponding proportion for LFPs was 20% (5 out of 25). We tested if Kruskal-Wallis gave different results as compared to ANOVA using Boschloo’s exact test, and found that neither the spike train data (p = 0.238) nor the LFPs (p = 0.378) were significantly different.

Since few of the sessions (within our 5% margin of error for false positives from α = 0.05) had a significant trial-to-trial variance in the spike trains, we concluded that we may perform pooled analysis of *d*_crit_ for the spike trains of trials across a single session. Boschloo’s exact test on the LFPs also demonstrated a non-significant deviation from the 5% margin of error (p = 0.413 for Kruskal-Wallis, p = 0.140 for ANOVA).

### Neural network structure is revealed by the DTW matrix

Instead of characterizing network connectivity with a single number (*d*_crit_), we used all the information available in the DTW matrix to determine if firing patterns were consistent, using non-parametric Mantel tests. We performed all pairwise comparisons for trials in the same rat to ask the question: given a single session, is there a difference in Pearson correlation coefficient of before vs. after depending on if we compare (1) before vs. after of the same trial to (2) before vs. after of different trials? We performed pairwise Mantel test comparisons among trials in the same session. We performed Kruskal-Wallis and Kolmogorov-Smirnov tests to determine if these distributions of the correlation coefficient and *p*-value were different (Supplemental Figures 2 and 3, Supplemental Table 3). Overall, we found that there was no significant difference trial-to-trial for the before vs. after DTW matrix correlation.

When we stratified by correctness (Supplemental Table 3), we did observe an effect for Kolmogorov-Smirnov tests on the Pearson correlation coefficient in 35.7% (5 out of 14) sessions. When we used *d*_crit_ alone, this proportion was 7.1% (1 out of 14). To evaluate if the KS-test/Mantel test combination was more powerful than the Kruskal-Wallis test on *d*_crit_, we performed a Boschloo exact test on the contingency table but this failed to reach statistical significance (p = 0.159). We concluded that there was mixed evidence for the efficacy of Mantel tests for before vs. after DTW matrices in determining if a rat made the correct or incorrect choice at the T-intersection.

### Neural network differences exist across multiple sessions for a single rat

Our second question was: when analyzing *d*_crit_, can we pool all the trials for a single rat, regardless of the day the recordings were taken? Using the Kruskal-Wallis test, we found that 70% (7 of 10) rats displayed significant (*p* < 0.05) day-to-day variance of *d*_crit_ in their spike trains, regardless of additional stratification by (1) correctness and (2) timing (Supplemental Table 1). The caveat was that we excluded six rats from the Yang study^16^ in this analysis, because we were unable to apply the Kruskal-Wallis test since recordings for those rats were only performed on a single day (dof = 0). Additionally, the three rats which did not display significant differences across sessions were 17914 and 18471 from the Ito study^14^ and rat A from Yang^16^. The same analysis was performed for LFPs, in which 80% (4 out of 5) rats (all rats except rat A from Yang^16^) displayed significant variance (Supplemental Table 2). For the LFPs, the five rats from Stout^15^ were excluded since the LFPs were not recorded for those rats. Boschloo’s exact test showed no difference between spike trains and LFPs (*p* = 0.922). We conclude that the resting cognitive state for each rat may be subject to change depending on the day that the recording is performed.

### Significant neural network heterogeneity exists between rats and studies

Our third question was: does *d*_crit_ depend on the specific rat? Using the Kruskal-Wallis test, we found that the Stout^15^ and Yang^16^ studies displayed significant (*p* < 0.001) rat-to-rat differences in their spike trains, whereas Ito^14^ was non-significant (*p* = 0.206). Ito^14^, however also had the fewest rats (*n* = 3) potentially leading to the insufficient statistical power of the Kruskal-Wallis test. The LFPs showed significant (*p* < 0.001) rat-to-rat variation in both Ito^14^ and Yang^16^. Pooling all rats, we confirmed (*p* < 0.001) that the specific study under consideration affects the value of *d*_crit_ for both spike trains and LFPs. We conclude that significant heterogeneities exist among studies, animals, and sessions. The statistical tests supporting this conclusion are summarized in Supplemental Tables 1 (spike trains) and 2 (LFPs).

### DTW distance is inconsistently correlated with physical electrode distance

Do correlations in firing among neurons reflect the underlying spatial distance between the neurons? We used both Welch’s t-test for unequal variances and the non-parametric Kolmogorov-Smirnov test (KS-test) to test for differences in the distribution of the DTW distance when grouped by the spatial distance of the associated electrodes. In all three studies, the *d*_crit_ between neurons recorded from the same electrode versus from different electrodes were statistically significantly different (*p* < 0.001) by both the t-test and KS-test (Supplemental Figures 4, 6-8); however, this classification was useless as a predictor or regressor^20^. For example, using *d_cutoff_* as a classifier of *d*_crit_ for determining if two neurons were recorded from the same electrode vs. from different electrodes, the receiver operator characteristic (ROC) curve is essentially no different from random guessing (Supplemental Figure 5), with area under the curve (AUC) of 0.51. If we stratify data by session (Supplemental Figures 6-8), however, the ROC curve and AUC vary for specific rats/sessions (Supplemental Figure 9), and in some cases demonstrate high predictive value (AUC).

For the Yang study^16^, we were able to obtain the exact geometry of the electrode. Therefore, we were able to stratify DTW distances by the physical distance of the neurons measured (Figure 3). For this stratification, we were unable to determine a consistent trend. Rats B and C demonstrate decreased DTW distance (increased synchronicity) in both neural spike trains and LFPs, whereas the remaining rats demonstrated slightly increasing DTW distance (decreased synchronicity) in both, except for rat S. For rat S, however, the large decrease in DTW distance seen in the spike trains may not be captured in the LFPs, because there was no zero-distance comparison for LFPs, whereas multiple spike trains recorded from the same electrode could be thought of as having zero distance.

**Figure 3:**
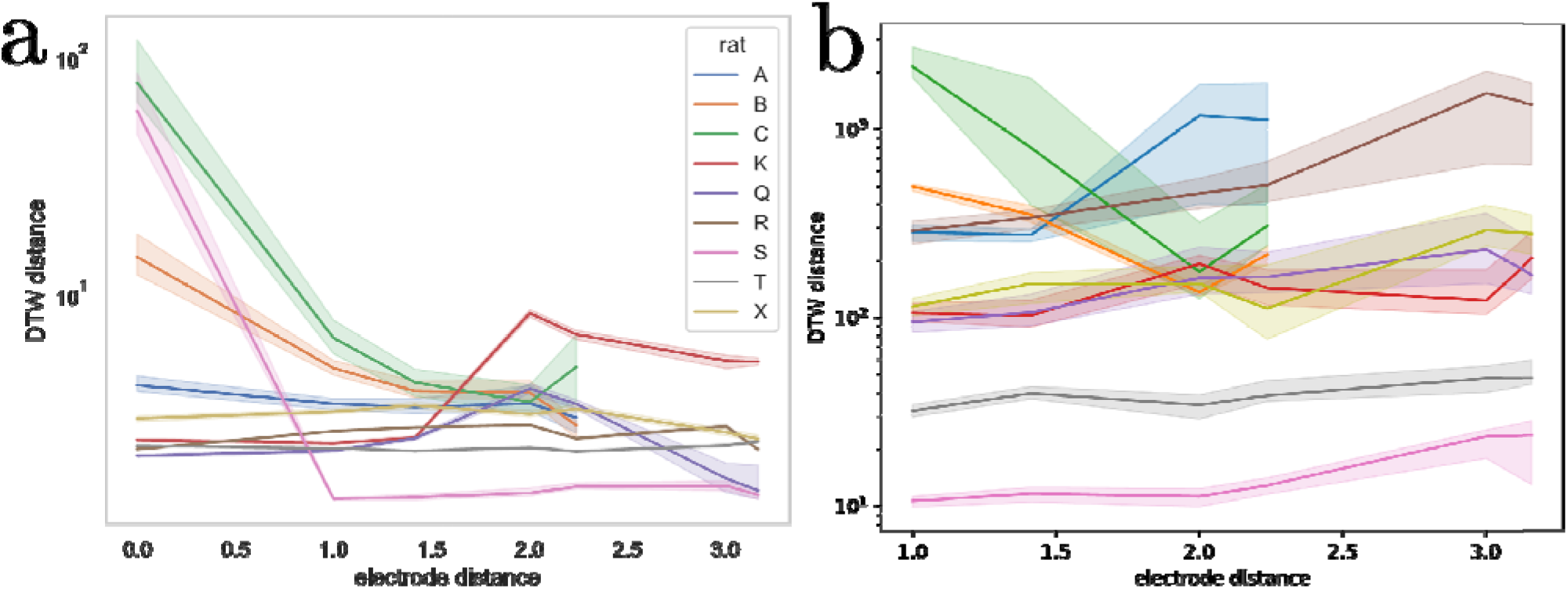
DTW distance for neuron spike trains (left) and LFPs (right) as a function of electrode distance (from Yang ^16^). The median DTW distance is plotted for each electrode distance, across all sessions for the indicated rat (rat A had 4 sessions, rat C had 3 sessions, and the remaining rats had a single session). 95% confidence intervals were estimated using 1,000 bootstrap samples. There is no consistent correlation between the DTW and electrode distance for either dataset, supporting the idea that there is no discernible spatial organization of neurons in the mPFC. Note that the DTW metric is different for the two datasets, with the distance for neuron spike trains reported as a time (ms) and the distance for LFPs reported as a voltage (V). Physical distance is reported in multiples of 0.25 mm.

## Discussion

We demonstrated that the DTW distance and the computed parameter *d*_crit_ captured some of the mPFC neural network firing dynamics for both the spike trains and LFPs, which were associated with T-maze task performance. We found weak evidence that the correctness of rat choice influences the firing dynamics (Supplemental Table 3). More importantly, we also found that significant heterogeneities exist among studies, animals, and sessions, as measured by DTW distance and *d*_crit_ To demonstrate meaningful associations between behavior (e.g., T-maze task performance) and neural network activities (in the mPFC), the data and computed results must be consistent. We first address the question of statistical consistency through a theoretical discussion of the experiments and the assumptions made in neuroelectrophysiology.

There are two major issues with attempting to measure a specific population of neurons. First, precise surgical implantation of electrodes is required; second, enough neurons must be sampled to capture overall neural network dynamics that can be consistently associated with certain behaviors. These two issues synergize at our current level of intracranial electrode technology, complicating fine measurements of neurons. Wheras the rat brain is far smaller than a human brain (ca. 1 cm^3^), it still contains an estimated 21 million neurons^21^. Reproducible surgical implantation in a specific area of the rat brain such as the medial prefrontal cortex is therefore highly dependent on the fine-motor skill of the researcher, and there is no guarantee that the same neurons or circuits will be sampled.

In fact, in Ito^14^ and Stout^15^ the electrodes were purposefully adjusted after each session so that a different set of neurons would be sampled. Due to the lack of spatial organization of the prefrontal cortex, we postulate that the surgical sampling of neurons and circuits in the prefrontal cortex is essentially random, and that electrode adjustment similarly resamples the neurons and circuits being recorded. The only way to counteract the statistical effects of random sampling is through the measurement of a sufficiently large fraction of the neurons in the brain area under consideration. In the Methods, we illustrate these issues with a simple mathematical illustration (*Reproducibility of electrode recordings*, Figure 1-D).

To provide a guide for researchers investigating the connection between neuroelectrophysiology and behavior, we asked what did each of the three studies do right and what could be improved? There are three experimental parameters we considered: *n_neurons_*(the number of neurons recorded); *n_network repeat_* (the number of times the same neural network is recorded); and *n_rats_*(the number of biological replicates). We contend that all three of these parameters must be relatively high to make meaningful conclusions, especially if one wishes to use more powerful parametric statistical tests over the non-parametric tests we used in this study.

To have any hope of identifying the specific neural network firing patterns which lead to decision-making in the T-maze, we must consistently identify the neurons responsible (*n_neurons_*), the circuits responsible (*n_network repeat_*), and demonstrate that the conclusions generalize across organisms (*n_rats_*). By this simple reasoning, we contend that the three recent studies we have examined here do not allow us to draw conclusions, regardless of data analysis method (DTW or otherwise). Stout^15^ recorded too few neurons (*n_neurons_*∼1), and changed the neurons recorded across sessions (*n_network repeat_* = 1) for *n_rats_* = 5. Ito^14^ recorded a moderate number of neurons (*n_neurons_* ∼20), but still changed the neurons recorded across sessions (*n_network repeat_* = 1), with *n_rats_* = 3. Yang^16^ recorded a moderate number of neurons (*n_neurons_* ∼20), and kept the same neurons recorded across sessions (*n_network repeat_* = 5), but only performed these replicates for *n_rats_* = 2 the other *n_rats_* = 7 had (*n_network repeat_* = 1). In short, larger electrodes (*n_neurons_*), more sessions per experimental condition (*n_network repeat_*), and more animals (*n_rats_*) are needed to achieve a robust and reproducible conclusion. Our recommendation for electrode size (*n_neurons_*) depends on the number of neural clusters or circuits one wishes to investigate (Figure 1-D), whereas *n_network repeat_*and *n_rats_* should be chosen according to traditional recommendations such that parametric statistical tests can be performed^22,23^.

We now turn to the efficacy of DTW in potentially uncovering connections between neuron electrical activity and behavior. In our analysis, we demonstrated that the DTW distance and the computed parameter *d*_crit_ captured some of the mPFC neural network firing dynamics for both the spike trains and LFPs. We found weak evidence that the correctness of rat choice influences the firing dynamics (Supplemental Table 3). We expect, however, that many more experiments are needed to confirm the classification of the correctness of a decision based on mPFC activity because our results indicated that whereas firing trials could be pooled across the same session, they could not be pooled across different sessions or studies (Supplemental Tables 1 and 2). In terms of spatial dependence of firing synchronicity, we found that for specific rats and sessions, the classification of DTW distances for recordings taken from the same vs. different electrodes was both statistically significant (Supplemental Figures 4, 6-8) and useful as a regressor (Supplemental Figure 9). For the Yang study^16^, the spatial dependence of DTW distances was highly dependent on the specific rat sampled (Figure 3), providing evidence of the lack of spatial organization of the prefrontal cortex.

The difficulties we encountered in analyzing spike trains and LFPs can be addressed through changes to the experimental design. The number of neurons sampled is an important characteristic to consider, especially if only a few neurons were recorded and some recordings represented only a single neuron (e.g., Stout^15^). If the mPFC is composed of separate neural networks that perform specific, modular tasks, it is unlikely that recording a low number of neurons will provide meaningful, interpretable results. Since firing dynamics were consistent across a single session, we posit that multiple recordings of a single neuron population are the best approach to characterizing network firing behavior. The DTW spatial dependence remained consistent across multiple sessions for rats A and C (Figure 3), therefore we are reasonably confident that we recorded the same population of neurons across the sessions. In contrast, because the electrodes were adjusted after each session, Ito^14^ and Stout^15^ only recorded a single session per neuron population that complicates the modelling we must perform. Is the additional variation across sessions in Yang^16^ due to changes in the resting cognitive state of the rats? For the data analyzed in this work, it was not possible for us to consistently distinguish between variance due to noisy measurements and variance due to rat behavioral changes. It may be necessary to perform many control sessions to characterize in adequate detail all of the resting cognitive states of a specific rat. Crucially, recording a single control session as Yang^16^ did for the seven other rats (B, K, Q, R, S, T, X) likely fails to detect variations in the resting state for a single rat. To complement the recording of many control sessions, recording additional rat behaviors may yield insight as to how the rat resting cognitive state influences the network firing dynamics.

Recent advances in imaging and brain modelling provide hope that we may one day understand the relationship between neuron firing and consciousness. Whole brain connectomes have been mapped for simple organisms (*C. elegans*^24^*, Ciona intestinalis* larvae^25^, and *Platynereis dumerilii*^26^), and recently all 3,000 neurons and 548,000 synapses of a fruit fly larva (*Drosophila melanogaster*) were mapped^27^. Although mammalian brains are orders of magnitude larger than insect brains, whole-brain connectomes for rats may not be out of reach in the coming decades. These connectomes are essential for understanding cognition, but they provide only a static, anatomic reference upon which neural physiology and pathophysiology must be built. Our study highlights the data requirements for the next steps in investigating the function and dynamics of biological neural networks.

## Methods and Materials

### Acquisition of additional rat recordings

Besides the data we used from the three published studies, new data were collected from three additional male Sprague-Dawley rats (Charles River Laboratories). They weighed 300–350 g when received and were named as rats A, B, and C. The experimental procedure of T-maze alternation and neural recording followed published protocol^16^. All animal care and surgical procedures were in accordance with the National Institutes of Health Guide for the Care and Use of Laboratory Animals and Penn State Hershey Animal Resources Program, and were reviewed and approved by the local IACUC.

### Dynamic time warping and statistical tests

Mathematical details on dynamic time warping, the Mantel test, and the Kolmogorov-Smirnov test are included in the Supplemental Methods.

### Estimated number of neurons in the rat medial prefrontal cortex

Our goal here is to provide a reasonable estimate for the number of neurons that one must record to reconstruct the dynamics of the entire mPFC. We used existing estimates of neuron and synaptic density in the rat prefrontal cortex as starting points for our modeling. We set the total number of neurons in the rat brain^21^ to *N_brain_* = 2.1 x 10^7^ with volume^28^ *V_brain_* = 2,500 mm^3^. We set the volume of the mPFC^28^ to *V_mPFC_* = 20 mm^3^. Assuming a uniform distribution of neuron count throughout the brain, the estimated number of neurons in the mPFC was 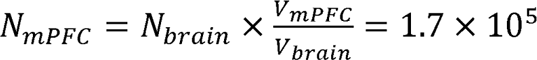.

### Reproducibility of electrode recordings

Here, we discuss the problem of reproducibility in neuroelectrophysiology. Because of the apparent lack of spatial organization in the mPFC, when electrodes are implanted, they uniformly sample the estimated *N_mPFC_*= 10^5^ neurons in the mPFC. Depending on how cooperative and synchronized neuron firing is, we assume that there is a certain number of neural clusters *n_cluster_* for which the measurement of a single neuron sufficiently captures the behavior of the entire cluster, and that measurement of these clusters correlates with rat behavior. We also assumed that sampling is done with replacement, i.e., *n_cluster_* << *N_mPFC._* For reproducibility, one must (1) record the same neural clusters across biological replicates and (2) identify or classify the neural clusters to show that findings in one animal generalize to other animals.

(1) The first issue of capturing a certain percentage of the neural clusters in the rat brain corresponds to a classical problem in combinatorial probability known as the coupon collection problem. The mathematical question is: given *c = n_cluster_* categories, what is the expected number of samples *S* which needs to be drawn from those categories so that all categories are represented at least once? In the case of equal probabilities for drawing each category,

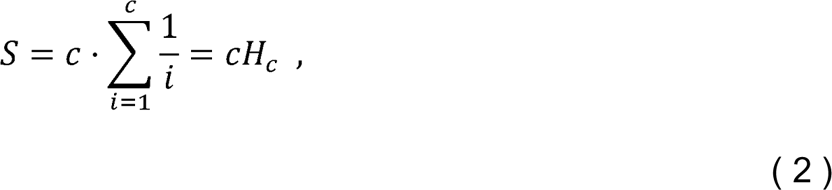

where *H_c_* is the *c^th^* harmonic number. Alternatively, we may require that a certain number *1 < k < c* out of all the categories is represented^29^, so that Eq. (2) is a special case of

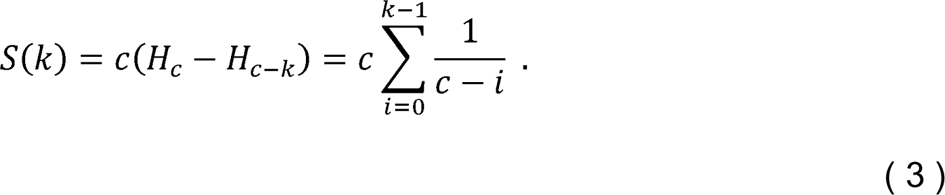

In both cases, *S ∼ c* log *c* asymptotically. We have plotted the expected number of clusters we may identify depending on the maximum number of neurons recorded from each of the studies (Figure 1-D). In practice, it is necessary to first determine the number of neural clusters that exist and to choose neuron sample size that can adequately sample all the clusters with high probability.

The second issue of identifying neural clusters across different animals relies on the researcher’s ability to determine which of the *n_cluster_*! possible matchings is the correct matching between any two rats. To reduce the number of possibilities, one must rely either on prior information (such as typical firing rates or patterns of certain clusters) or on other measurements of network behavior that are invariant under permutations of nodes in the graph.

### Computational analysis and plotting

All computational analysis was carried out in Python 3.10 and R 4.1, on an M1 Max MacBook Pro with 64 GB RAM. For statistical tests: Shapiro-Wilk, Kruskal-Wallis, and ANOVA were carried out using pingouin 0.5.3^30^, Mantel tests^31,32^ were carried out using scikit-bio 0.5.7^31^, and Kolmogorov-Smirnov and Boschloo exact tests were carried out using SciPy 1.8.1^33^. We implemented dynamic time warping^19^ using Cython 0.29.32^34^. We used the DataFrame structure from Pandas 1.5.1^35^ to organize our data, and we used seaborn 0.12.1^36^ and Matplotlib 3.5.2^37^ for plotting. We used NumPy 1.23.3^38,39^ for numerical array operations. We used ggpubr 0.6.0^40^ for the balloon plots.

## Acknowledgements

We thank Hiroshi T. Ito, May-Britt Moser, John J. Stout, and Amy L. Griffin for supplying rat neuroelectrophysiology recordings and precise experimental details for our data analysis.

We acknowledge support from the National Institutes of Health (NIH) 1R35 GM134864 (N.V.D.), 1RF1 AG071675 (Y.Y., N.V.D., R.B.M.), 1R01 AT012053 (N.V.D.), the National Science Foundation 2210963 (N.V.D.), and the Passan Foundation (N.V.D.).

## Author contributions (CRediT statement)

C.M.S. Conceptualization, methodology, software, validation, formal analysis, investigation, data curation, writing – original draft, writing – review & editing, visualization. J.W. Conceptualization, methodology, software, formal analysis, writing – review & editing.

R.B.M. Conceptualization, methodology, validation, resources, writing – review & editing, supervision, project administration, funding acquisition.

Y.Y. Conceptualization, methodology, validation, investigation, resources, data curation, writing – review & editing, supervision, project administration, funding acquisition.

N.V.D. Conceptualization, methodology, software, validation, formal analysis, resources, writing – review & editing, supervision, project administration, funding acquisition.

## Competing interests

We confirm that we have no financial or other interests related to this work which could affect or have the perception of affecting our objectivity or the content of this article.

## Code and Data Availability

All code and data which are necessary to reproduce the results in this work are posted on Zenodo (https://doi.org/10.5281/zenodo.7933346).

## Abbreviations

(NEP): Neuroelectrophysiology
(DTW): dynamic time warping
(mPFC): medial prefrontal cortex
(KS-test): Kolmogorov-Smirnov test
(ROC): receiver operating characteristic
(AUC): area under the curve

## Notes

https://doi.org/10.5281/zenodo.7933346

### Competing Interest Statement

The authors have declared no competing interest.

### Summary of Updates

Reorganized abstract and discussion to discuss theoretical results before statistical results; expanded introduction to better motivate the use of dynamic time warping; updated language in manuscript and Supplementary material ("event" -> "trial", "run" -> "session"); standardized author affiliations (all authors were from the same institution)

