## Supplemental Tables for "Quantifying network behavior in the rat prefrontal cortex: a reproducibility crisis"

Supplemental Table 1: Statistical tests for neuron spike trains.

| **Categorical variable** | **Null hypothesis (H_0_)** | **Alternative hypothesis (H_A_)** | **Test** | **Additional restrictions/stratification** | **Fraction of tests with p < 0.05 (number out of total)** |
| --- | --- | --- | --- | --- | --- |
| None | The $d_{\text{crit}}$ are normally distributed for each session. | The $d_{\text{crit}}$ are NOT normally distributed for each session. | Shapiro-Wilk | None | 0.967 (60 out of 62 sessions) |
|  |  |  |  | correctness = true | 0.967 (60 out of 62 sessions) |
|  |  |  |  | correctness = false | 0.967 (60 out of 62 sessions) |
|  |  |  |  | timing = before | 0.967 (60 out of 62 sessions) |
|  |  |  |  | timing = after | 0.967 (60 out of 62 sessions) |
|  |  |  |  | timing = before AND correctness = true | 0.967 (60 out of 62 sessions) |
|  |  |  |  | timing = before AND correctness = false | 0.967 (60 out of 62 sessions) |
|  |  |  |  | timing = after AND correctness = true | 0.967 (60 out of 62 sessions) |
| Trials in a single session | The $d_{\text{crit}}$ for each trial does not vary across a session. | At least one trial in the session has a significantly different $d_{\text{crit}}$ from the others. | Kruskal-Wallis | None | 0.048 (3 out of 62 sessions) |
|  |  |  |  | correctness = true | 0.048 (3 out of 62 sessions) |
|  |  |  |  | correctness = false | 0.048 (3 out of 62 sessions) |
|  |  |  |  | timing = before | 0.048 (3 out of 62 sessions) |
|  |  |  |  | timing = after | 0.048 (3 out of 62 sessions) |
|  |  |  |  | timing = before AND correctness = true | 0.048 (3 out of 62 sessions) |
|  |  |  |  | timing = before AND correctness = false | 0.048 (3 out of 62 sessions) |
|  |  |  |  | timing = after AND correctness = true | 0.048 (3 out of 62 sessions) |
|  |  |  |  | timing = after AND correctness = false | 0.048 (3 out of 62 sessions) |
|  |  |  | ANOVA | None | 0.112 (7 out of 62 sessions) |
|  |  |  |  | correctness = true | 0.112 (7 out of 62 sessions) |
|  |  |  |  | correctness = false | 0.112 (7 out of 62 sessions) |
|  |  |  |  | timing = before | 0.112 (7 out of 62 sessions) |
|  |  |  |  | timing = after | 0.112 (7 out of 62 sessions) |
|  |  |  |  | timing = before AND correctness = true | 0.112 (7 out of 62 sessions) |
|  |  |  |  | timing = before AND correctness = false | 0.112 (7 out of 62 sessions) |
|  |  |  |  | timing = after AND correctness = true | 0.112 (7 out of 62 sessions) |
|  |  |  |  | timing = after AND correctness = false | 0.112 (7 out of 62 sessions) |
| Trials across all sessions for a rat | The $d_{\text{crit}}$ for each trial does not vary across all sessions for a rat. | At least one trial across all sessions for the rat has a significantly different $d_{\text{crit}}$ from the others. | Kruskal-Wallis | None | 0.700 (7 out of 10 rats) |
|  |  |  |  | correctness = true | 0.700 (7 out of 10 rats) |
|  |  |  |  | correctness = false | 0.700 (7 out of 10 rats) |
|  |  |  |  | timing = before | 0.700 (7 out of 10 rats) |
|  |  |  |  | timing = after | 0.700 (7 out of 10 rats) |
|  |  |  |  | timing = before AND correctness = true | 0.700 (7 out of 10 rats) |
|  |  |  |  | timing = before AND correctness = false | 0.700 (7 out of 10 rats) |
|  |  |  |  | timing = after AND correctness = true | 0.700 (7 out of 10 rats) |
| Trials across all trials for all rats in the study | The population of $d_{\text{crit}}$ sampled is independent of rat in the study. | At least one trial across all rats in the study has a significantly different $d_{\text{crit}}$ from the others. | Kruskal-Wallis | None | 0.667 (2 out of 3 studies) |
|  |  |  |  | correctness = true | 0.667 (2 out of 3 studies) |
|  |  |  |  | correctness = false | 0.667 (2 out of 3 studies) |
|  |  |  |  | timing = before | 0.667 (2 out of 3 studies) |
|  |  |  |  | timing = after | 0.667 (2 out of 3 studies) |
|  |  |  |  | timing = before AND correctness = true | 0.667 (2 out of 3 studies) |
|  |  |  |  | timing = before AND correctness = false | 0.667 (2 out of 3 studies) |
|  |  |  |  | timing = after AND correctness = true | 0.667 (2 out of 3 studies) |

Supplemental Table 2: Statistical tests for local field potentials.

| **Categorical variable** | **Null hypothesis (H_0_)** | **Alternative hypothesis (H_A_)** | **Test** | **Additional restrictions** | **Fraction of tests with p < 0.05 (number out of total)** |
| --- | --- | --- | --- | --- | --- |
| None | The $d_{\text{crit}}$ are normally distributed for each session. | The $d_{\text{crit}}$ are NOT normally distributed for each session. | Shapiro-Wilk | None | 0.800 (20 out of 25 sessions) |
|  |  |  |  | correctness = true | 0.800 (20 out of 25 sessions) |
|  |  |  |  | correctness = false | 0.800 (20 out of 25 sessions) |
|  |  |  |  | timing = before | 0.800 (20 out of 25 sessions) |
|  |  |  |  | timing = after | 0.800 (20 out of 25 sessions) |
|  |  |  |  | timing = before AND correctness = true | 0.800 (20 out of 25 sessions) |
|  |  |  |  | timing = before AND correctness = false | 0.800 (20 out of 25 sessions) |
|  |  |  |  | timing = after AND correctness = true | 0.800 (20 out of 25 sessions) |
| Trials in a single session | The $d_{\text{crit}}$ for each trial does not vary across a session. | At least one trial in the session has a significantly different $d_{\text{crit}}$ from the others. | Kruskal-Wallis | None | 0.120 (3 out of 25 sessions) |
|  |  |  |  | correctness = true | 0.120 (3 out of 25 sessions) |
|  |  |  |  | correctness = false | 0.120 (3 out of 25 sessions) |
|  |  |  |  | timing = before | 0.120 (3 out of 25 sessions) |
|  |  |  |  | timing = after | 0.120 (3 out of 25 sessions) |
|  |  |  |  | timing = before AND correctness = true | 0.120 (3 out of 25 sessions) |
|  |  |  |  | timing = before AND correctness = false | 0.120 (3 out of 25 sessions) |
|  |  |  |  | timing = after AND correctness = true | 0.120 (3 out of 25 sessions) |
|  |  |  |  | timing = after AND correctness = false | 0.120 (3 out of 25 sessions) |
|  |  |  | ANOVA | None | 0.200 (5 out of 25 sessions) |
|  |  |  |  | correctness = true | 0.200 (5 out of 25 sessions) |
|  |  |  |  | correctness = false | 0.200 (5 out of 25 sessions) |
|  |  |  |  | timing = before | 0.200 (5 out of 25 sessions) |
|  |  |  |  | timing = after | 0.200 (5 out of 25 sessions) |
|  |  |  |  | timing = before AND correctness = true | 0.200 (5 out of 25 sessions) |
|  |  |  |  | timing = before AND correctness = false | 0.200 (5 out of 25 sessions) |
|  |  |  |  | timing = after AND correctness = true | 0.200 (5 out of 25 sessions) |
|  |  |  |  | timing = after AND correctness = false | 0.200 (5 out of 25 sessions) |
| Trials across all sessions for a rat | The $d_{\text{crit}}$ for each trial does not vary across all sessions for a rat. | At least one trial across all sessions for the rat has a significantly different $d_{\text{crit}}$ from the others. | Kruskal-Wallis | None | 0.800 (4 out of 5 rats) |
|  |  |  |  | correctness = true | 0.800 (4 out of 5 rats) |
|  |  |  |  | correctness = false | 0.800 (4 out of 5 rats) |
|  |  |  |  | timing = before | 0.800 (4 out of 5 rats) |
|  |  |  |  | timing = after | 0.800 (4 out of 5 rats) |
|  |  |  |  | timing = before AND correctness = true | 0.800 (4 out of 5 rats) |
|  |  |  |  | timing = before AND correctness = false | 0.800 (4 out of 5 rats) |
|  |  |  |  | timing = after AND correctness = true | 0.800 (4 out of 5 rats) |
| Trials across a study | The population of $d_{\text{crit}}$ sampled is independent of rat in the study. | At least one trial across all rats in the study has a significantly different $d_{\text{crit}}$ from the others. | Kruskal-Wallis | None | 1.000 (2 out of 2 studies) |
|  |  |  |  | correctness = true | 1.000 (2 out of 2 studies) |
|  |  |  |  | correctness = false | 1.000 (2 out of 2 studies) |
|  |  |  |  | timing = before | 1.000 (2 out of 2 studies) |
|  |  |  |  | timing = after | 1.000 (2 out of 2 studies) |
|  |  |  |  | timing = before AND correctness = true | 1.000 (2 out of 2 studies) |
|  |  |  |  | timing = before AND correctness = false | 1.000 (2 out of 2 studies) |
|  |  |  |  | timing = after AND correctness = true | 1.000 (2 out of 2 studies) |

Supplemental Table 3: Tests on changes in before vs after spike train DTW matrix correlation.

| **Categorical variable** | **Null hypothesis (H_0_)** | **Alternative hypothesis (H_A_)** | **Dependent variable** | **Test** | **Fraction of tests with p < 0.05 (number out of total)** |
| --- | --- | --- | --- | --- | --- |
| Same vs different trial in a single session | There is no difference for Mantel tests of before vs after depending on if the before and after matrices are taken from the same vs different trials in a single session. | There is a difference in Mantel test results depending on if the before and after matrices are taken from the same vs different trials in a single session. | Pearson correlation coefficient (r) | Kruskal-Wallis | 0.045 (2 out of 44) |
|  |  |  |  | Kolmogorov-Smirnov | 0.045 (2 out of 44) |
|  |  |  | Mantel test p-value | Kruskal-Wallis | 0.069 (3 out of 43) |
|  |  |  |  | Kolmogorov-Smirnov | 0.045 (2 out of 44) |
| Correct vs incorrect trials in a single session | There is no difference for Mantel tests of before vs after depending on if the before and after matrices are taken from the correct vs incorrect trials in a single session. | There is a difference in Mantel test results depending on if the before and after matrices are taken from the correct vs incorrect trials in a single session. | Pearson correlation coefficient (r) | Kruskal-Wallis | 0.000 (0 out of 13) |
|  |  |  |  | Kolmogorov-Smirnov | 0.357 (5 out of 14) |
|  |  |  | Mantel test p-value | Kruskal-Wallis | 0.000 (0 out of 9) |
|  |  |  |  | Kolmogorov-Smirnov | 0.214 (3 out of 14) |
| Correct vs incorrect trials in a single session | $d_{\text{crit}}$ is independent of correctness of trials in a single session. | $d_{\text{crit}}$ depends on correctness of trials in a single session. | $d_{\text{crit}}$ | Kruskal-Wallis | 0.071 (1 out of 14) |
