## Supplemental Figures for "Quantifying network behavior in the rat prefrontal cortex: a reproducibility crisis"

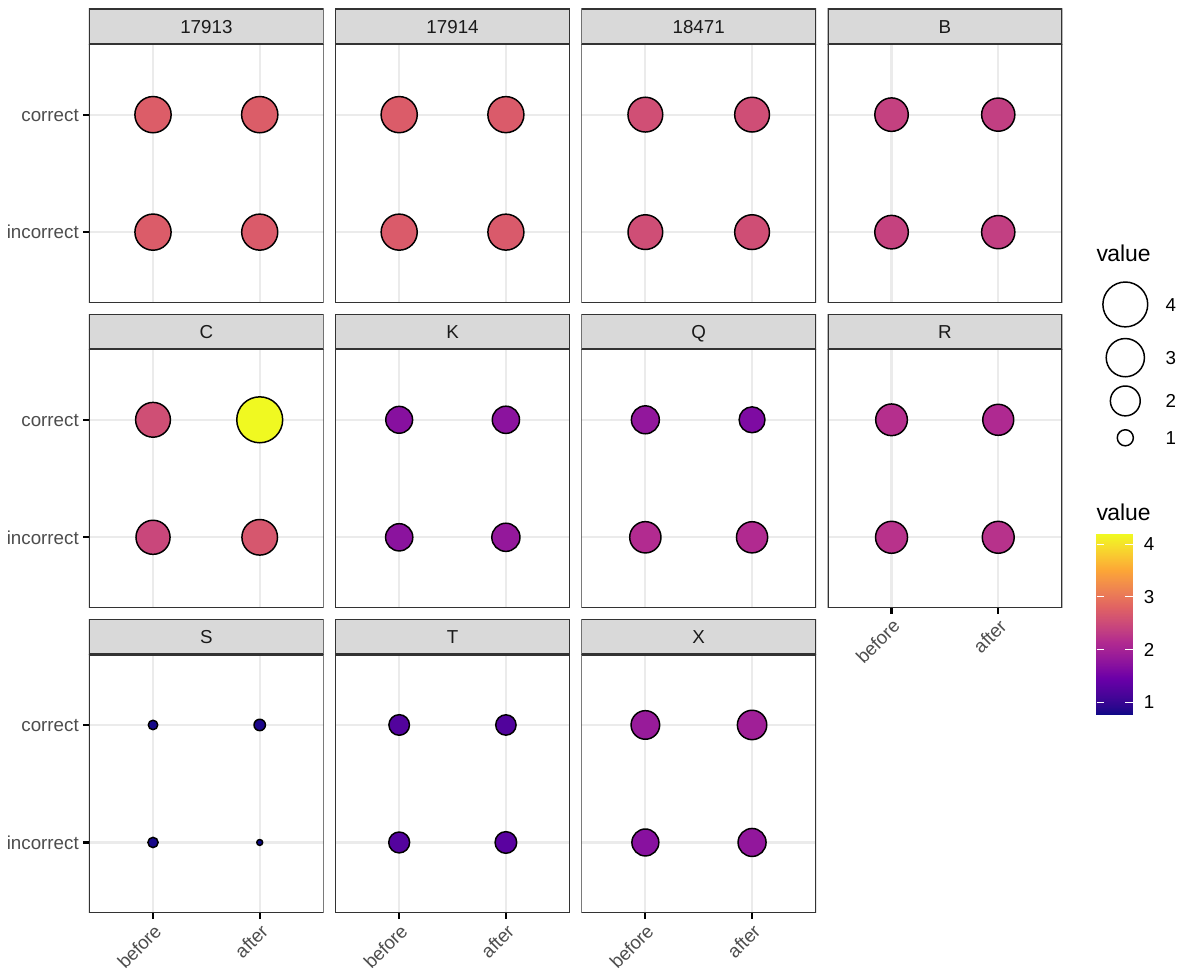

Supplemental Figure 1: Balloon plot of local field potential $d_{\text{crit}}$, grouped by rat, correctness of T-arm choice, and timing with respect to T-arm choice. Each box represents a single rat. The x-axis represents timing (before vs after the rat visits the T-maze intersection) and the y-axis represents correctness of T-arm taking (True vs False). The size and color of each marker represents $\log_{10} d_{\text{crit}}$. For rats with multiple trials, we took the mean of $\log_{10} d_{\text{crit}}$. The scale of $\log_{10} d_{\text{crit}}$ is larger than that for the spike trains, visualized in Figure 1. The balloon plot visually appears more homogeneous. Local field potentials were not recorded in the Stout 2020 study.

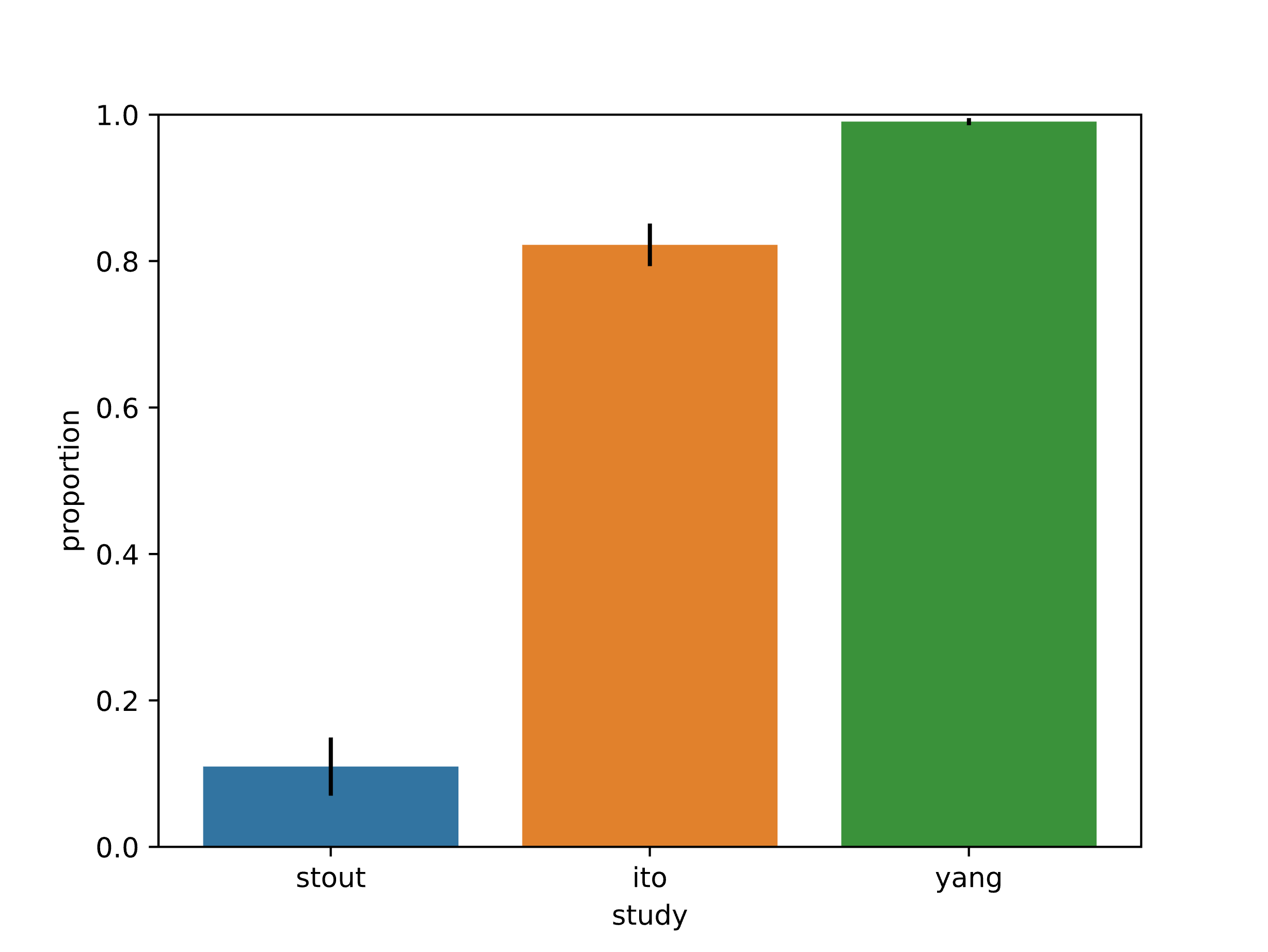

Supplemental Figure 2: The proportion of Mantel tests which were significant (*p* < 0.05), stratified by study. A significant Mantel test indicates significant correlation between the two spike train DTW matrices which were compared, and therefore an indication of the consistency of recordings. We plotted 95% confidence intervals using the Agresti-Coull interval. As expected, the Yang 2022 study was the most consistent, since the electrodes were fixed across trials and sessions. In contrast, recordings from Stout 2020 were limited in the number of neurons recorded, and the electrodes were also not fixed, resulting in a low proportion of significant Mantel tests.

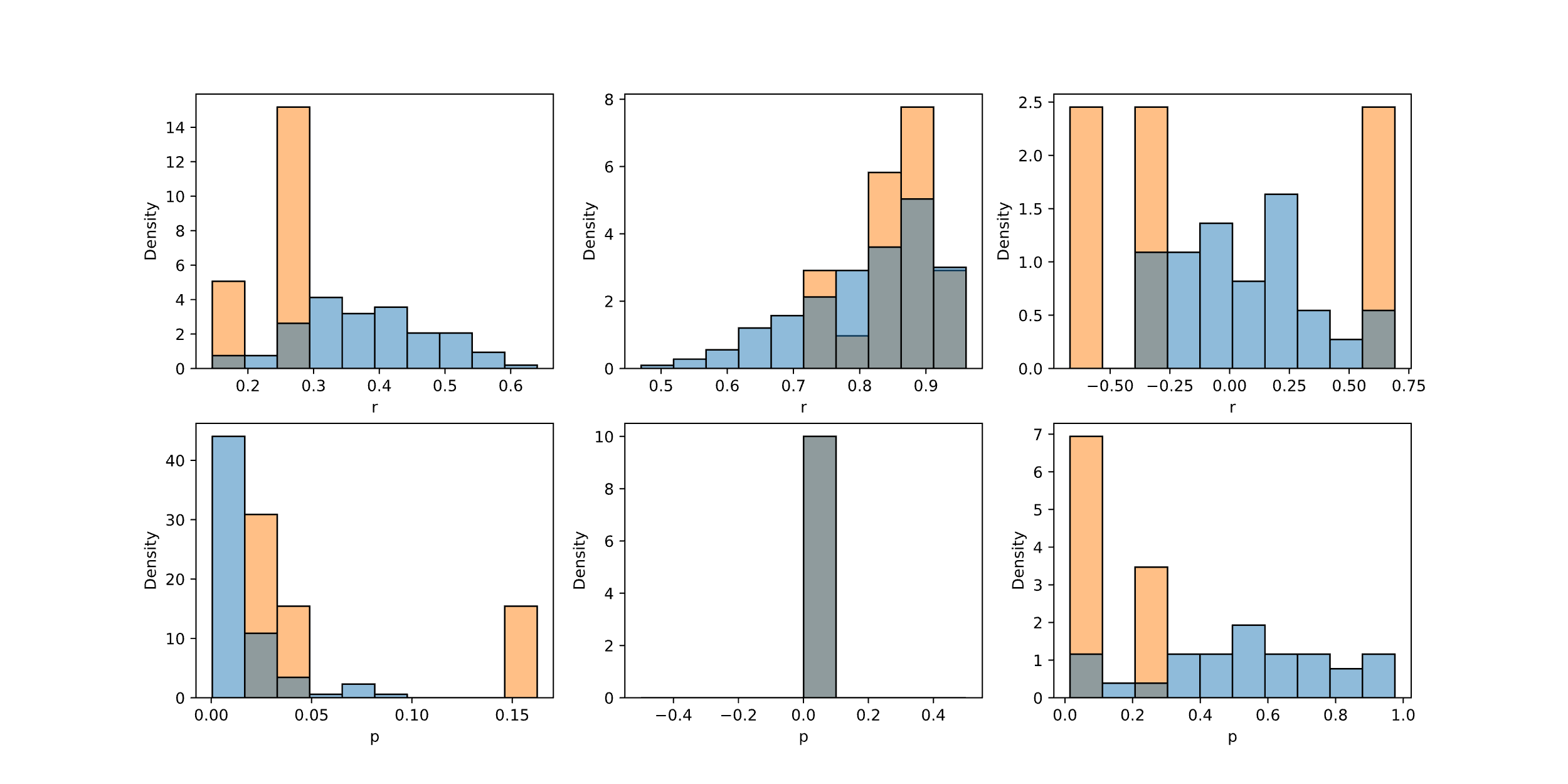

Supplemental Figure 3: Mantel test correlation and p-value distribution for trials across three sessions, stratified by comparison. From left to right, the columns represent sessions: (1) 17914, Feb 24, 2013; (2) Q, Mar 11, 2015; and (3) Groot, Mar 13, 2018. The top row shows Pearson correlation coefficient distributions, and the bottom row shows p-value distributions. The blue represents before vs after comparisons for different trials in the same session, whereas the orange represents before vs after comparisons for the same trial. Using Kruskal-Wallis tests, (1) and (2) had different distributions of correlation coefficient (*p* < 0.05), while (1) and (3) had different distributions of p-values (*p* < 0.05).

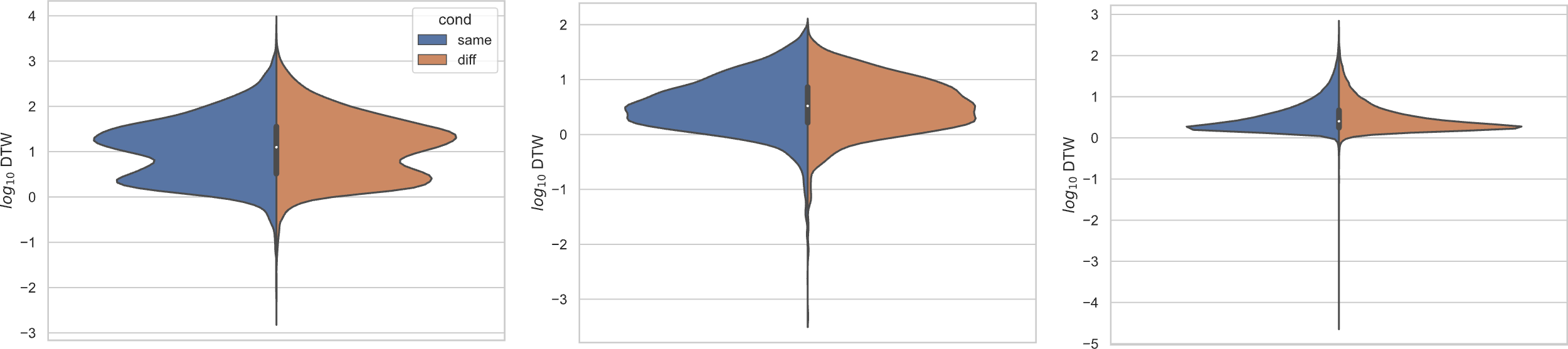

Supplemental Figure 4: Comparing distributions for DTW matrix entries for neuron spike trains, depending on if the neurons compared were from the same vs different electrode. Left: Ito 2018 (n=25,542 same, n=173,368 diff), middle: Stout 2020 (n=8,462 same, n=10,186 diff), right: Yang 2022 (n=125,580 same, n=763,588 diff). For each study, Kolmogorov-Smirnov tests demonstrated significant (*p* < 0.001) differences between same vs different electrode data.

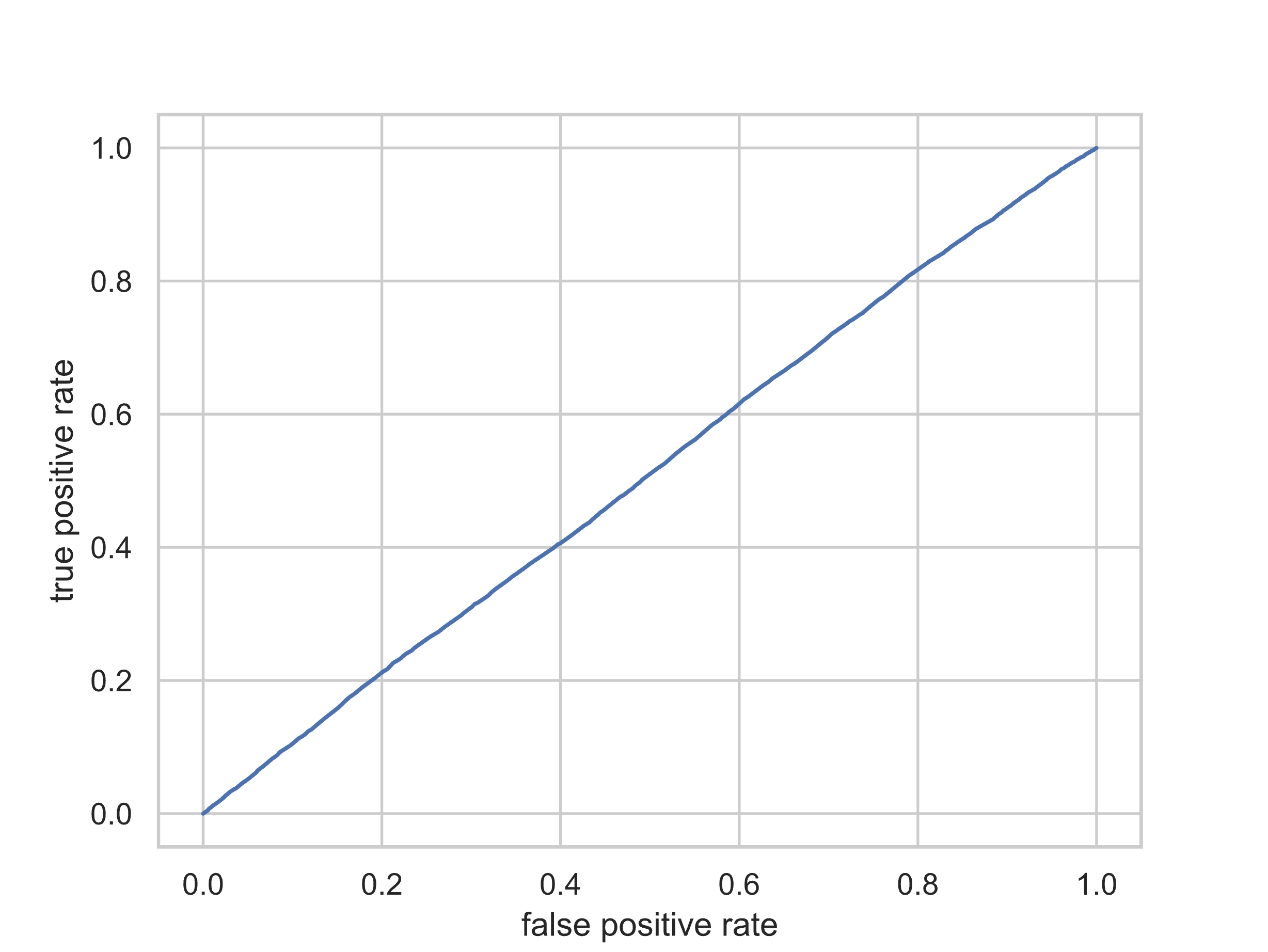

Supplemental Figure 5: ROC curve for spatial data from Ito 2018, using cutoffs for the DTW matrix value as a predictor for the categorical variable same vs different electrode. The AUC is 0.51, indicating limited predictive value.

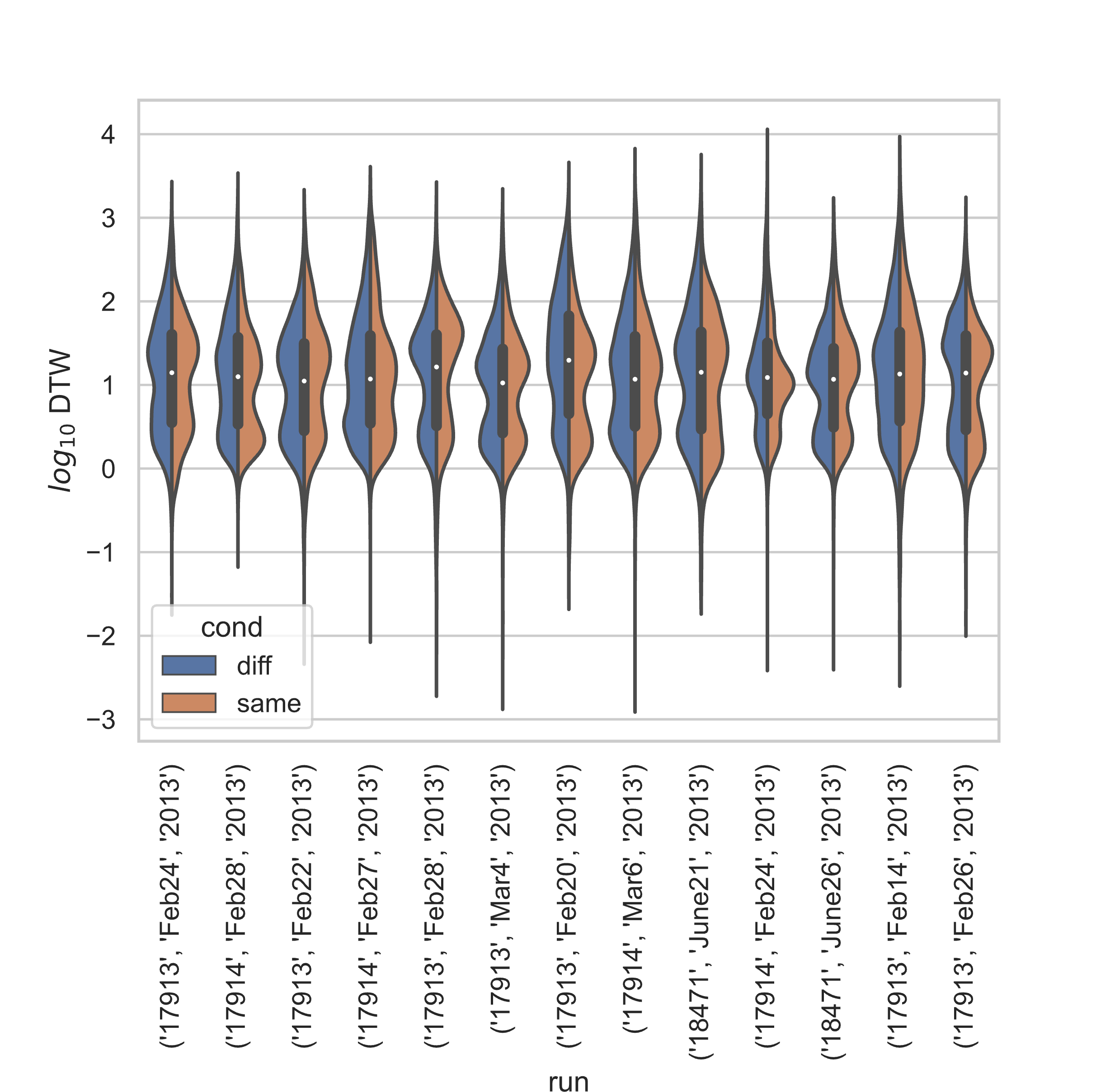

Supplemental Figure 6: Stratified comparison of distributions for DTW matrix entries for neuron spike trains, for Ito 2018, depending on if the neurons compared were from the same vs different electrode. For all 13 sessions, Kolmogorov-Smirnov tests demonstrated significant (*p* < 0.001) differences between the two populations.

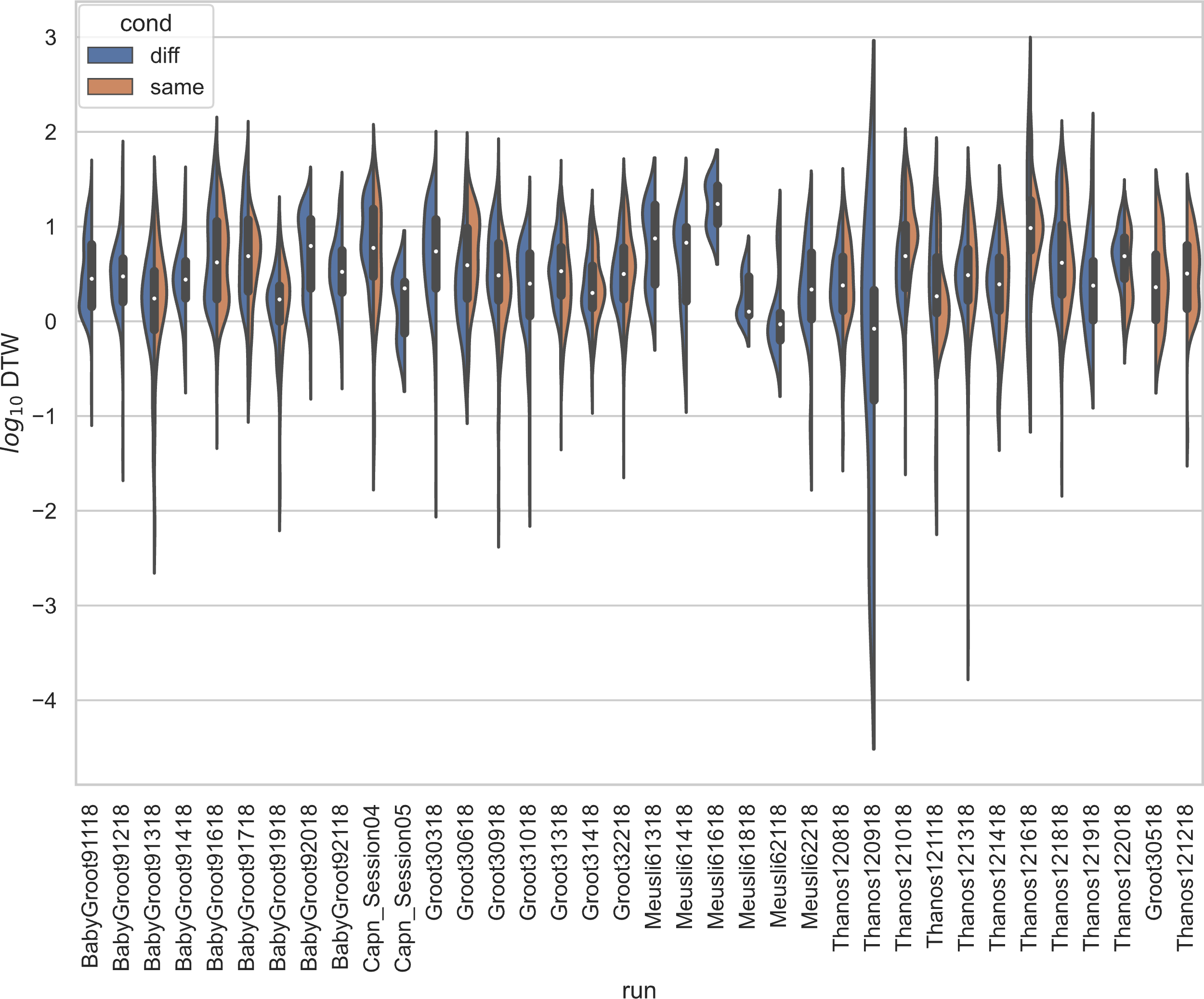

Supplemental Figure 7: Stratified comparison of distributions for DTW matrix entries for neuron spike trains, for Stout 2020, depending on if the neurons compared were from the same vs different electrode. For all 34 sessions with non-empty groups of same vs diff, Kolmogorov-Smirnov tests demonstrated significant (*p* < 0.001, with the exception of session Thanos121618, for which *p* = 0.007) differences between the two populations.

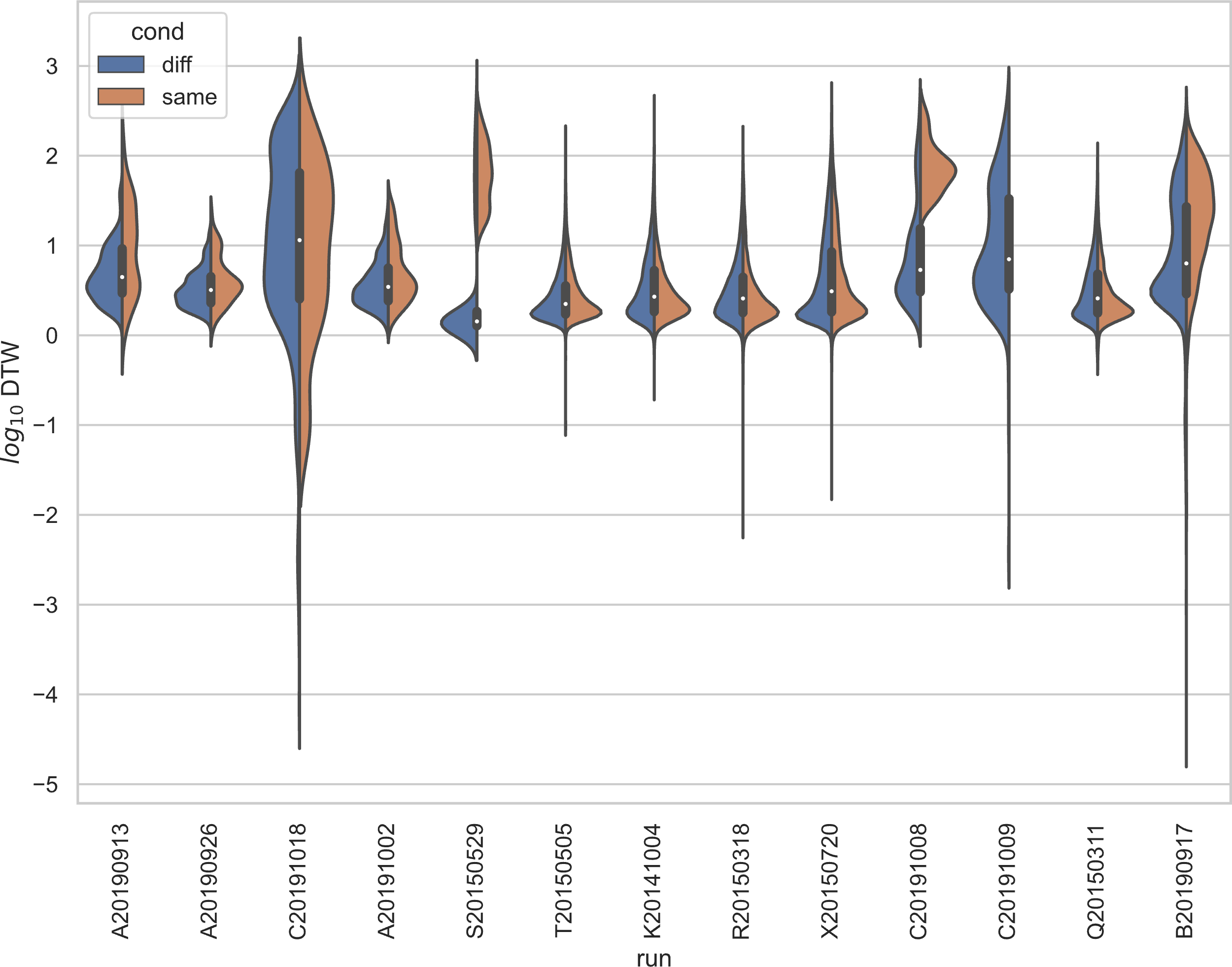

Supplemental Figure 8: Stratified comparison of distributions for DTW matrix entries for neuron spike trains, for Yang 2022, depending on if the neurons compared were from the same vs different electrode. For all 13 sessions, Kolmogorov-Smirnov tests demonstrated significant (*p* < 0.001) differences between the two populations.

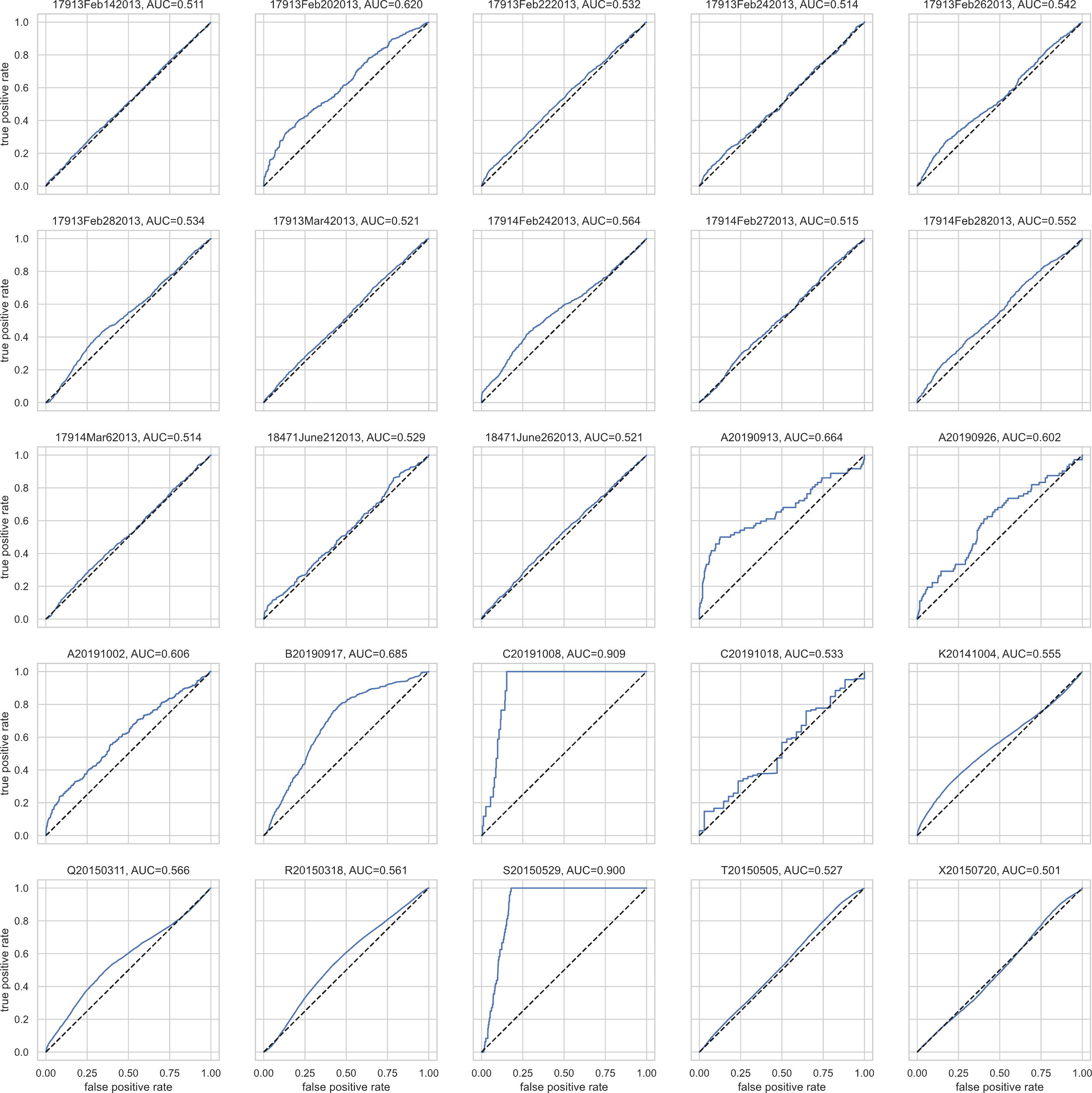

Supplemental Figure 9: ROCs for each of the Ito 2018 and Yang 2022 sessions, using a certain value of DTW distance as a cutoff for classifying the reading as from the same vs different electrodes. We omit Stout here because the number of DTW entries per matrix was not comparable. There is significant variability in AUC across the sessions. The dotted line indicates random guessing; when the ROC dips below the dotted line, DTW matrix entries are consistently being classified into the wrong class.
