## Supplemental Methods for "Quantifying network behavior in the rat prefrontal cortex: a reproducibility crisis"

*Dynamic time warping*

Dynamic time warping (DTW) is a method for time series and sequence alignment^1^. Given two sequences of data $x_{i}=x\left( u_{i} \right)$and $y_{j}=y\left( v_{j} \right)$ with lengths $m$ and $n$ respectively, DTW seeks to find a monotonically increasing function $w: u_{i}\to v_{j}$, such that $w\left( u_{1} \right)=v_{1}$, $w\left( u_{m} \right)=v_{n}$, such that each $u_{i}$ is paired with a $v_{j}$ and vice versa, and such that a measure of sequence distance $d(x, y)$ is minimized. This problem possesses an optimal substructure and therefore may be treated using dynamic programming for an efficient solution.

When the sequence distance is a sum of costs for each pair of values

$$d\left( x,y \right)=\sum_{i} d_{s}(x\left( u_{i} \right),y(w(u_{i}))$$

( 1 )

we can express DTW as an algorithm on a matrix running in $O\left( mn \right)$ time.

Consider an alignment of $x$ and $y$. The last symbols must be mapped to each other $x_{m}\leftrightarrow y_{n}$, and must be included in the total distance. There are three possibilities for how this alignment came about. First, $y_{n}$ could be mapped to a previous $x_{i}, i<m$; second, $y_{n}$ and $x_{m}$ could be matched to each other, separately from the shorter sequences $\{y_{1},\cdots,y_{n-1}\}$ and $\{x_{1},\cdots,x_{m-1}\}$; third, $x_{m}$ could be matched to a previous $y_{j}$, $j<n$. Denoting the distance of the alignment of the first $i$ entries of $x$ and the first $j$ entries of $y$ as $M(i,j)$, we have the recursion relation

$$M(m,n)=d_{s}\left( x_{m},y_{m} \right)+\min\left\{ M\left( m-1,n \right),M\left( m-1,n-1 \right),M\left( m,n-1 \right) \right\} .$$

( 2 )

The DTW method is shown in Algorithm 1.

We performed dynamic time warping on both the neuron spike trains and local field potentials (LFPs). Since the spike trains were all-or-none trials and each entry of $x$ and $y$ represented an action potential at time $t_{i}$ and $t_{j}$ respectively, we used the distance function

$$d_{spike}\left( x\left( t_{i} \right),y\left( t_{j} \right) \right)=\left| t_{i}-t_{j} \right| .$$

( 3 )

For the local field potential, the measured quantity was the voltage, and we used the distance function

$$d_{LFP}\left( x\left( t_{i} \right),y\left( t_{j} \right) \right)=\left| x\left( t_{i} \right)-y\left( t_{j} \right) \right| .$$

( 4 )

For each pair of neuron spike trains for a given rat and each pair of LFP tracings, we computed the overall dynamic time warping distance according to Eq. ( 1 ), forming a distance matrix. One major caveat is that the DTW distance does not satisfy the triangle inequality and therefore is not a metric. However, DTW distances may be projected onto lower dimensions through principal coordinates analysis, classical multidimensional scaling, or clustering techniques for analysis.

*Mantel test*

To compare the neuron DTW distance matrices for spike trains and LFPs, we employed the Mantel test^2,3^, a non-parametric test of matrix similarity. The idea of the Mantel test is to mitigate correlations among matrix entries by permuting the columns of one of the matrices. For a pair of matrices $(M,N)$, the Mantel test computes a Pearson correlation coefficient $\rho$

$$\rho\left( M,N \right)=\frac{\text{cov}\left( M_{ij},N_{ij} \right)}{\sigma_{M}\sigma_{N}}=\frac{\sum_{ij} (M_{ij}-\bar{M})(N_{ij}-\bar{N})}{\sqrt{\sum_{ij} \left( M_{ij}-\bar{M} \right)^{2}}\sqrt{\sum_{ij} \left( N_{ij}-\bar{N} \right)^{2}}} ;$$

( 5 )

$\rho(M,N)$ is then compared to the distribution of $\rho_{perm}\left( M,N \right)$, in which Eq. ( 5 ) is computed for all permutations of the columns (or rows) of $M$ (or $N$) while keeping the other matrix fixed. For large matrices, the permutations are sampled from the set of all permutations.

The location of $\rho\left( M,N \right)$ with respect to all the $\rho_{perm}\left( M,N \right)$ gives an estimate of the similarity of the matrices.

*Kolmogorov-Smirnov test*

To compare DTW matrices between rats, we used the Kolmogorov-Smirnov (KS) test^4,5^ to compare distributions of the relative percentiles of the Pearson correlation coefficient computed from the Mantel test. The KS test is a parameter-free test which determines if samples drawn from two populations represent different underlying probability distributions.

First, the empirical distribution function ($P_{1}$ and $P_{2}$) of each population is estimated based on the samples,

$$P_{k}\left( x \right)=\sum_{i} \delta\left( x-s_{k,i} \right) ,$$

( 6 )

where $\delta(x)$ is the Dirac delta function and $s_{k,i}$ is the $i^{th}$ sample from distribution $k$. Second, the empirical cumulative distribution function ($\text{F}_{1}$ and $\text{F}_{2}$) for each population is estimated as

$$F_{k}\left( x \right)=\int_{-\infty}^{x} P_{k}\left( t \right) \text{dt}\text{ .}$$

( 7 )

Finally, the KS statistic is computed:

$$D=\sup_{x} \left| F_{1}\left( x \right)-F_{2}\left( x \right) \right| .$$

( 8 )

For a confidence level $\alpha$, the null hypothesis is rejected (i.e. the underlying probability distributions are different) if

$$D>\sqrt{-\log\left( \frac{\alpha}{2} \right)\cdot\left[ \frac{1+\frac{m}{n}}{2m} \right]} .$$

( 9 )

where $m$ and $n$ are the sample sizes.
